# The role of miR-31-5p in the development of intervertebral disc degeneration and its therapeutic potential

**DOI:** 10.1101/2020.09.16.299982

**Authors:** Yong Zhou, Jiqing Su, Mingsi Deng, Wei Zhang, Dongbiao Liu, Zhengguang Wang

## Abstract

Intervertebral disc degeneration (IDD) refers to the abnormal response of cell-mediated progressive structural failure. In order to understand the molecular mechanism of the maintenance and destruction of the intervertebral disc, new IDD treatment methods are developed. Here, we first analyzed the key regulators of IDD through miRNA microarrays. The cell structure and morphology were discovered by Histological and radiographic. Then, the level of miR-31-5p was disclosed by qRT-PCR. The association between miR-31-5p and SDF-1/CXCR7 axis was discovered by 3′-Untranslated region (UTR) cloning and luciferase assay. The apoptosis of cells under different treatments was disclosed by Flow cytometer. The cell proliferation was discovered by EdU assay. Finally, the protein levels of SDF-1, CXCR7, ADAMTS-5, Col II, Aggrecan and MMP13 were discovered by Western blot. The results show that miR-31-5p is a key regulator of IDD and its level is down-regulated in IDD. Overexpression of miR-31-5p facilitates NP cell proliferation, inhibits apoptosis, facilitates ECM formation and inhibits the level of matrix degrading enzymes in NP cells. The SDF-1/CXCR7 axis is the direct target of miR-31-5p. miR-31-5p acts on IDD by regulating SDF-1/CXCR7. In vitro experiments further verified that the up-regulation of miR-31-5p prevented the development of IDD. In conclusion, overexpression of miR-31-5p can inhibit IDD by regulating SDF-1/CXCR7.

## Introduction

Epidemiological statistics show that with the development of economy and society, the prevalence of low back pain worldwide has been increasing year by year, which has surpassed spinal cord injury and has become one of the diseases with the highest disability rate [1]. The main root cause of low back pain is intervertebral disc degeneration (IDD) [2]. IDD refers to the abnormal response of cell-mediated progressive structural failure [3]. IDD is a complex pathophysiological process, which is caused by the interaction between the nucleus pulposus cells, extracellular matrix and the biomechanics of the intervertebral disc, thus forming a closed vicious circle process, and continue to promote the development of IDD [4]. The gelatinous nucleus pulposus (NP) of the ID is anatomically surrounded by anal ring fibrosis. The occurrence of degeneration will lead to the imbalance of the internal environment of the intervertebral disc, loss of tissue hydration, inflammation, and loss of extracellular matrix, which will lead to a decrease in the height of the intervertebral disc, destruction of the annulus fibrosus structure, and a gradual loss of normal physiological structure and function [5]. IDD is a chronic process of degradation and destruction of extracellular matrix protein (ECM). The degradation products of ECM may trigger or further promote the inflammatory response interrelated to intervertebral disc degeneration and low back pain [6]. Therefore, inhibiting the phenotypic abnormality of NP cells is the crucial to preventing the progression of IDD.

The pathogenesis of IDD is interrelated to many biological and genetic regulators [7, 8], but it has recently been discovered that MicroRNAs (miRNAs) are important regulators of the development of IDD. miRNAs regulate protein level by binding to the 3’untranslated region of mRNAs, and then play a part in a variety of physiological and pathological processes [9]. miRNAs are key regulators of a variety of cellular physiological processes, including proliferation, differentiation, apoptosis, survival and morphogenesis. A large number of studies have manifested that miRNAs are widely present in tissues and organs, and play a regulatory part in the occurrence and development of various diseases [10]. Abnormal miRNA level is present in various musculoskeletal diseases, such as osteoporosis and rheumatoid arthritis [11–13]. More and more evidence support that miRNAs play a part in the process of causing IDD [14–16]. The study found that miR-494, miR-27a, miR-155, miR-93, miR-146a, miR-377, miR-100, miR-21, miR-10, miR-146a and other miRNA s have been reported to be interrelated to the development of IDD [17]. However, the association between miR-31-5p and IDD is still less studied.

miR-31-5p is one of the most popular candidate genes [18]. Emerging evidence shows that miR-31-5p is an oncogenic or tumor suppressor gene in different types of tumors, and it is also a useful clinical prognostic biomarker [19]. In IDD tissues, the level of miR-31-5p, miR-124 A and miR-127-5p is generally down-regulated [20]. It is interrelated to the role of ECM synthesis in hypertrophic scar formation [21]. This study confirmed the abnormal regulation of miR-31-5p in NP tissues of IDD patients. Subsequently, we obtained degenerated human NP cells from IDD patients to study the pathways through which miR-31-5p functions in IDD.

## Materials and Methods

### Patient samples

NP samples were taken from 82 underwent discectomy patients with degenerative disc disease (57.6 ± 5.3 years in total). The indications for surgery are failure of conservative treatment and progressive neurological deficits, such as progressive motor weakness or cauda equina syndrome. Here we exclude patients with lumbar spine stenosis, ankylosing spondylitis, isthmus or degenerative spondylolisthesis or diffuse idiopathic skeletal hypertrophy. A total of 91 patients were recruited to be used as the age- and sex-matched controls, which involved collecting NP tissue samples taken from fresh traumatic lumbar fracture as the result of anterior decompressive surgery. Before surgery, these patients underwent a routine lumbar spine MRI scan. According to Pfirrmann classification, the degree of intervertebral disc degeneration was graded. The research protocol has been approved by the Ethics Committee of The Third Xiangya Hospital Affiliated to Central South University, and the written informed consent of each participant has been obtained.

### IDD model

In this study, an IDD model was established in mice (12 weeks old) by AF needle puncture [22, 23]. The selected mouse is degenerated tail disc, because the tail disc is most similar to human lumbar disc [24–26]. Ketamine (100 mg/kg) was selected as the anesthetic for mice in the operation group, and they were injected intraperitoneally. After the general anesthesia effect was achieved, the mouse was fixed in the left side position and executed the model operation. A sagittal small skin incision was performed from Co6 to Co8 to help locate the disc position for needle insertion in the tail. Subsequently, Co6–Co7 coccygeal discs were punctured using a syringe needle. The syringe needle was inserted into Co6–Co7 disc along vertical direction and then rotated in the axial direction by 180° and held for 10 s. Using a 31-G needle, puncture the NP through the AF parallel to the end plate through the NP, and then insert it into the 1.5 mm disc to decompress the nucleus. The other parts remain unchanged as the comparison part. For treatment experiments, intervertebral discs were harvested from WT mice at 6 and 12 weeks after surgery and then studied.

### Primary nucleus pulposus (NP) monolayer culture

The NP tissues were washed 3 times with phosphate-buffered saline (PBS; Gibco, Grand Island, NY), minced into small fragments and digested in 0.25% (w/v) trypsin (Gibco) and 0.2% (w/v) type Π collagenase (Gibco) and then placed in PBS for approximately 3 hours at 37 °C in a gyratory shaker. Cells were filtered through a 70 μm mesh filter (BD, Franklin Lakes, NJ). Primary NP cells were cultured with growth medium (Dulbecco’sModified Eagle’s Medium and Ham’s F-12 Nutrient Mixture (DMEM-F12; Gibco), 20% (v/v) fetal bovine serum (FBS; Gibco), 50 U/mL penicillin, and 50 μg/mL streptomycin (Gibco) in 100mm culture dishes in a 5% (v/v) CO_2_ incubator. The cells were passaged at approximately 80% (v/v) confluence using trypsin and subcultured in a 60mm culture dish (2.5×10^5^ cells/well). Cells that had been passaged no more than twice were used in the subsequent experiments.

### MiR-31-5p NP generation and injection

Use Silencer^®^ siRNA labeling kit (#AM1636) to transfect cultured primary human NP cells with Cy3 labeled or unlabeled miR-31-5p mimics or inhibitors. Use Lipofectamine RNAiMAX Transfection Reagent (Invitrogen) at 50 nM. To inhibit the level of SDF-1/CXCR7, cells were transfected with siRNA (Thermo Scientific Dharmacon®) using Lipofectamine 3000 (Invitrogen) according to the manufacturer’s instructions. For treatment experiments, we divided 48 male mice (12-week-old C57BL/6) (Bar Harbor Jackson Laboratory, USA) that underwent IDD surgery into four groups (12 mice in each group). In the Cy3 simulation control group NP treatment group or Cy3-miR-31-5p simulation NP treatment group, mice were locally injected with 20 μL Cy3-miR simulation control NP or 20 μL Cy3-miR-31-5p simulation NP on the 1,7,14 day. In order to determine the transfection efficiency of the Cy3-labeled miR-31-5p mimic/inhibitor or its negative control, IVIS 200 imaging system (Xenogen, Calper Life Science, MA, USA) was applied for in vivo fluorescence imaging, and at different times Perform histological examination at points after each group injection (24, 48 and 72 h). In the 6th and 12th weeks after the operation, the intervertebral discs were collected for histological and radiological evaluation of each group.

### Histological and radiographic evaluation

The intervertebral discs of the mice were fixed in 10% neutral formalin buffer for 1 week, and then soaked in EDTA decalcification solution for decalcification. Then it was embedded and sectioned. Then the histological images were analyzed by hematoxylin, eosin and saffron O-type green staining using Olympus BX51 microscope (Olympus Center Valley, Pennsylvania, USA). Based on a literature review of research on intervertebral disc degeneration, an improved histological grading system was developed [27–33]. Specific operation and evaluation refer to the reference article given.

### qRT-PCR

Total RNA was extracted from the transfected cell lines using Trizol reagent (Takara, Japan). One-step PrimeScript miRNA cDNA Synthesis Kit (Takara) was applied for reverse transcription. We applied SYBR Geen Realtime PCR Master Mix (Takara) to perform qRT-PCR and synthesize data on the ABI 7300 system (ABI). In addition, U6 snRNA were applied as internal controls.

### CpG island prediction and Bisulfite sequencing PCR (BSP)

The promoter region was predicted by using Promoter Inspector prediction software (http://www.genomatix.ed). The CpG prediction algorithm was applied to predict the CpG islands associated with the promoter. Genomic DNA was isolated from NP by Qiagen DNeasy Blood and Tissue Kit, and then placed in bisulfite. Then it was amplified with BSP primers and cloned into pGEMT Easy vector (Promega, Wisconsin, USA). The samples are then sequenced and the data is analyzed by BIQ analyzer.

### Microarray analyses

First, the Trizol method was employed to extract total RNA from NP cells preserved with a final concentration of 1 mg·ml-1. Then the mi RNA isolation kit (Ambion) of mir Vana TM was employed to purify the mi RNA part of the total RNA, and finally the extracted mi RNA samples were analyzed on the chip, using the human mi RNA chip (v.12.0) of Agilent Company for analysis. Hybridization. Data processing was performed by GeneSpring GX v12.1 software package (Agilent Technologies).

### Transfection

We applied miR-31-5pmimic, miR-31-5p inhibitor, non-targeting siRNA control (si-NC) and miRNA control (RiboBio Inc., Guangzhou, China) according to the instructions of the Lipofectamine™ 3000 kit (Invitrogen, Carlsbad, CA) transfected into Cy3 labeled or unlabeled NP cells.

### Luciferase assay

The cells were planted in a 24-well plate at 10^5^ cells/well and cultivated for 24 h. According to the instructions of the detection kit (Promega), the cells were transfected with the luciferase reporter gene level plasmid, and the miR-31-5p mimic and the control group were given at the same time. After 24 h, the luciferase activity was measured.

### Flow cytometer (FCM)

Cell apoptosis was disclosed using FITC-Annexin V and ethidium iodide (PI, 556547, BD biosciences). For the operation method, refer to the kit instructions and calculated the percentage of apoptosis according to the fluorescence intensity.

### EdU analysis

After processing the cells according to the treatment conditions and time of each group, 50 μmol/L Edu (Sigma-Aldrich) medium was replenished to each dish. The cells were cultured in a 37°C, 5% CO_2_ incubator for 2 h, and rinsed with PBS twice. The cells were fixed with 4% paraformaldehyde, 0.2% glycine was replenished, and rinsed with PBS for 5 min. The membrane was ruptured by 0.5% Triton-100 for 10 minutes, rinsed with PBS, and 100 μL Apollo staining reaction solution was replenished to each well. It was incubated on a shaker at room temperature in the dark for 30 minutes, rinsed with 0.5% Triton-100, and then rinsed with 100 μL methanol and PBS. It was stained with DAPI for 20 min, then rinsed with PBS, and the results were observed under a fluorescence microscope.

### Fluorescence in situ hybridization

Locked nucleic acid (LNA) probes complementary to miR-31-5p are labeled with 5’and 3’-digoxigenin (Exiqon, Woburn, MA, USA). The NP tissue of IDD patients is employed for FISH detection. This section was taken out, and then the gene break probe was dropped. Then, it was placed on the hybridization instrument at 75°C for 10 min, 42°C overnight. It was taken out the next day, and rinsed at room temperature for 5 minutes and 72°C for 3 minutes. It was dried and DAPI was replenished dropwise to cover the slide. The fluorescence microscope (Olympus IX-81; Olympus, Tokyo, Japan) was employed to read and take the image. The intensity of miR-31-5p staining was scored from 0 to 4 based on no staining [34]. Then analyzed the miR-31-5p cells in three representative high-power fields of a single sample.

### Western blot

Collect cells in the logarithmic growth phase and seed them in a petri dish with a diameter of 60 mm. After 24 h of cell treatment, cells of each group were collected and employed RIPA lysate. The protein concentration of each group of cells was disclosed by BCA method, and 6% to 12% SDS-PAGE electrophoresis was executed with 20 μg/well of protein. The proteins separated by electrophoresis are transferred to the PVDF membrane. After incubating the first antibody, the second antibody anti-rabbit immunoglobulin G (ab99697) was incubated at room temperature for 1 h, and developed with ECL reagent (Thermo Fisher Scientific, Inc.). Use Image J software (National Institutes of Health) to analyze the gray value of each protein band to determine the optical density.

The primary antibody information is as follows: anti-Col II antibody (Abcam: ab34712), anti-agroglycan antibody (ab36861), anti-ADAMTS-5 antibody (ab41037), anti-MMP13 antibody (ab39012), anti-SDF-1 antibody (ab155090), anti-CXCR7 antibody (ab72100), anti-β-actin (ab150301).

### Cellular immunofluorescence

The cells were seeded on glass slides in a 6-well culture plate, transfected the next day, and 4% paraformaldehyde was employed to fix the cells on the glass slides 48 h later. Then it was permeabilized with 0.5% Triton X-100 for 20 minutes, blocked with normal goat serum for 30 minutes, and the primary antibody was dropped and incubated overnight at 4°C. These cells were incubated with FITC-labeled secondary antibody in the dark for 1 h at room temperature. The nuclei were counterstained with DAPI, rinsed, mounted and observed under a fluorescence microscope.

The primary antibody information is as follows: anti-Col II antibody (Abcam: ab34712), anti-agroglycan antibody (ab36861), anti-ADAMTS-5 antibody (ab41037), anti-MMP13 antibody (ab39012). The secondary antibody is goat anti-rabbit IgG (Abcam: ab150077).

### Immunofluorescence and TUNEL staining of tissue sections

The frozen part of the mouse dish was fixed with 4% paraformaldehyde. The tissue sections were taken out, and after deparaffinization and hydration, they were fixed in pre-cooled 40 g/L paraformaldehyde for 5 min. It was treated with proteinase K for 10 min, immersed in a buffer solution containing 5% hydrogen peroxide for 20 min to block the endogenous peroxidase activity. TUNEL (Invitrogen) reaction mixture was replenished and reacted at 37 °C for 1 h. POD conversion solution was replenished and reacted at 37 °C for 30 min. Then, DAB was developed and the apoptosis was observed under the microscope.

### Statistical analysis

All data were analyzed using SPSS 19.0 software (SPSS Inc., Chicago, Illinois, USA), and the experiment was repeated 3 times. The result is expressed as x±SD. The comparison between the two groups was executed using an independent sample t test. Multiple group comparisons were performed through one-way analysis of variance. *P*<0.05 was accounted significant.

## Result

### miR-31-5p declined in NP tissues of IDD patients

The occurrence of IDD is closely interrelated to the dysregulation of interrelated miRNAs[17]. In order to study the part of miRNAs in IDD, we disclosed miRNAs by microarray (Figure 1A). Then, these significantly dysregulated miRNAs were subjected to unsupervised cluster analysis to differentiate IDD patients from controls (Figure 1B and 1C). From Figure 1B and 1C, it can be observed that miR-31-5p has a significant imbalance. Therefore, we chose miR-31-5p for further research. We used further qRT-PCR experiments to discover miR-31-5p level in human nucleus pulposus tissues and nucleus pulposus cells. The results demonstrated that compared to the control group, the miR-31-5p in the nucleus pulposus tissue and nucleus pulposus cells of the IDD group was significantly reduced (Figure 1D and 1E, *P* < 0.001). We confirmed this conclusion through further fluorescence in situ hybridization experiments (Figure 1F). In order to probe the upstream mechanism of miR-31-5p down-regulation in NP, CpG islands in the miR-31-5p promoter region were predicted (Figure 1G). As we can see, the methylation status of IDD group was significantly higher than that of control group (Figure 1H, *P* < 0.001). From the above results, it was revealed that the level of miR-31-5p in the NP tissue of IDD patients declined and the methylation status increased.

**Figure. 1.**
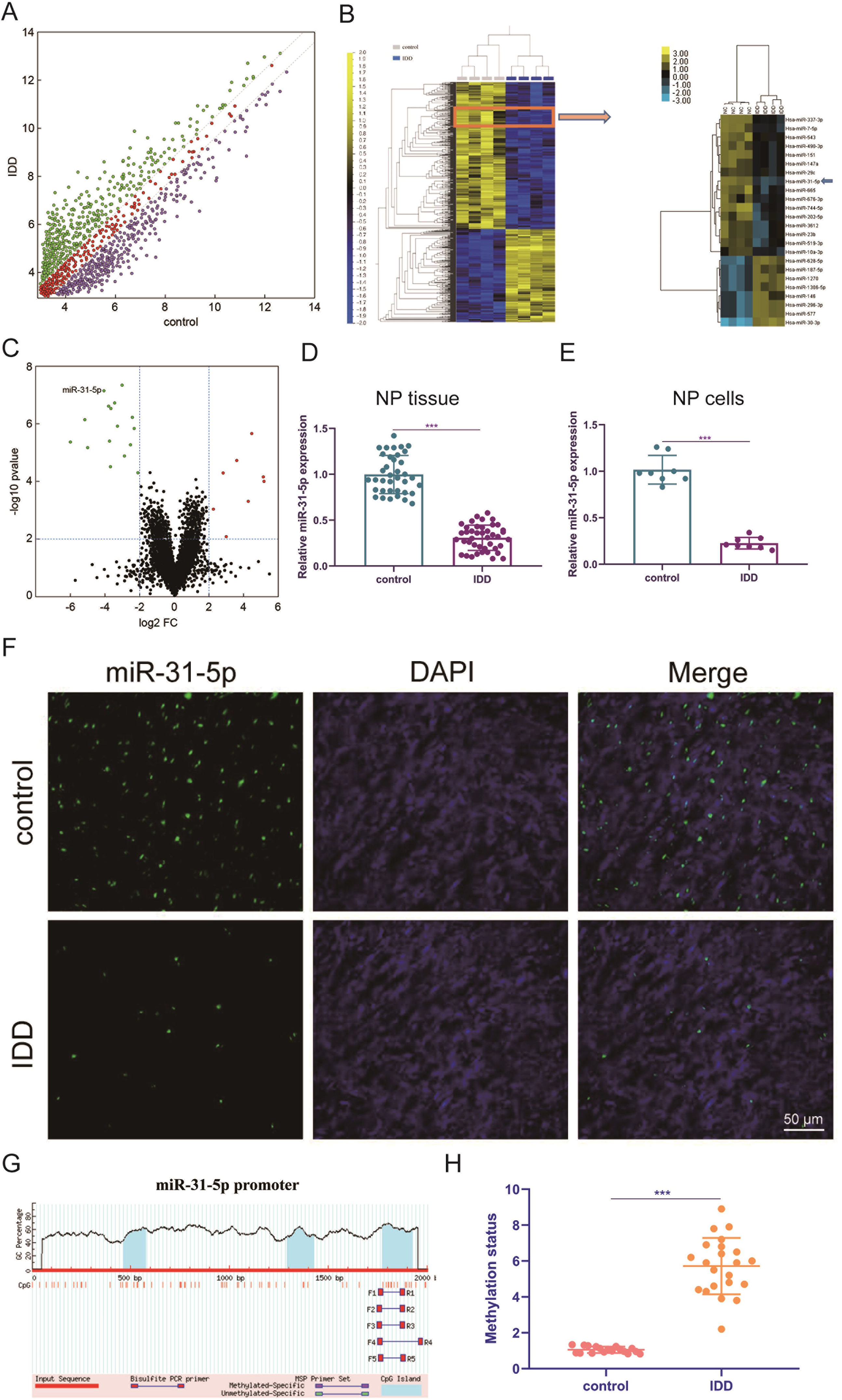
miR-31-5p declined in NP tissues of IDD patients. (A) Represents the scatter plot of miRNA expression profiles between IDD patients and the control group (green dots indicate more than two-fold increase; purple dots indicate more than two-fold down). (B) Describes the heat maps of 24 differentially expressed miRNAs. (C) The volcano graph represents the difference in miRNA expression levels between IDD patients and controls. The y-axis is the negative P value after adjusting Log10, and the x-axis represents the multiple change of Log2. The red dots on the right represent up-regulated miRNAs, and the green dots on the left represent down-regulated miRNAs. miR-31-5p is indicated. (D) The miR-31-5p level in Human nucleus pulposus tissue was disclosed through qRT-PCR. (E) The miR-31-5p level in Human nucleus pulposus cells was valued through qRT-PCR. (F) FISH analysis was executed on IDD patients and the control group. Scale bar = 50μm. (G) Methylation of miR-31-5p promoter region. (H) Methylation status of IDD patients and controls. n = 3. IDD, Intervertebral disc degeneration; miR: microRNA; FC: fold change; DAPI: 4’,6-diamidino-2-phenylindole. \*\*\**P <* 0.001.

### The effect of miR-31-5p overexpression or silence on the phenotype of NP cells

Previous studies have demonstrated that miR-31-5p is closely interrelated to the occurrence of IDD [35]. To further probe the part of miR-31-5p in IDD, the miR-31-5p mimic or inhibitor was transfected into primary human NP cells. It was demonstrated that the transfection efficiency of Cy3-labeled miRNA was disclosed (Figure 2A). We further studied the effects of miR-31-5p overexpression or silencing on NP cell proliferation, apoptosis, ECM formation and matrix-degrading enzymes. The results of EdU demonstrated that compared with miR-31-5p inhibitor, the upregulation of miR-31-5p level promoted NP cell proliferation (Figure 2B). In terms of apoptosis, upregulation of miR-31-5p level restrained NP cell apoptosis (Figure 2C). We further disclosed the function of miR-31-5p levels on anabolic/catabolism markers through the function gain and loss of function studies. It was demonstrated that the levels of Col II and Aggrecan increased in primary human NP cells transfected with miR-31-5p mimics. In primary human NP cells transfected with miR-31-5p inhibitor, the levels of ADAMTS-5 and MMP13 increased (Figure 2D). We further verified this function by immunofluorescence (Figure 2E and 2F). Overall, the data signifies that the overexpression of miR-31-5p facilitates the synthesis and proliferation of NP cell matrix.

**Figure.2.**
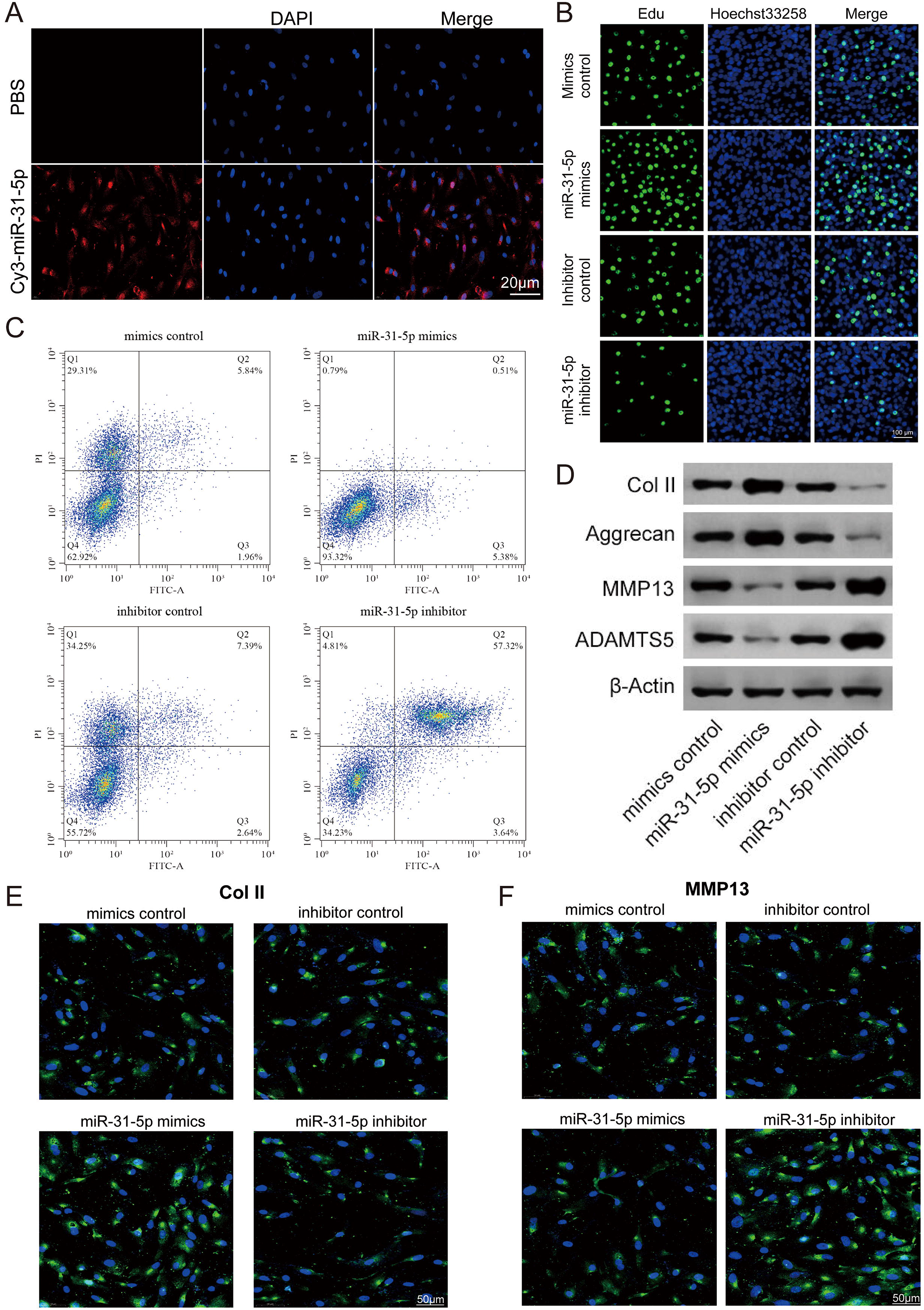
The effect of miR-31-5p overexpression or silence on the phenotype of NP cells. (A) Cy3 was employed to detect miR-31-5p transfected and cultured NP cells, scale bar = 20μm. (B) Analyze the proliferation of NP cells transfected with different treatments by EdU, scale bar = 100μm. (C) The apoptosis of NP cells was analyzed by FCM. (D) The level of MMP13, Col II, Aggrecan and ADAMT5 was valued through western blot. (E, F) The Col II and MMP13 levels were disclosed through immunofluorescence. N=3. IDD, Intervertebral disc degeneration; miR: microRNA; PBS: phosphate buffer saline; DAPI: 4’,6-diamidino-2-phenylindole; FITC: Fluorescein Isothiocyanate; Col II: Type II collagen; MMP: matrix metalloprotein; ADAMTS: a disintegrin-like and metalloproteinase with thrombospondin motifs; NP: Nucleus pulposus; EdU: 5-Ethynyl-2′-deoxyuridine.

### The relevance of miR-31-5p to SDF-1/CXCR7 axis

We performed a gene ontology (GO) analysis on the dysregulated mRNA. Our results demonstrated that in the biological process, the GO term of the down-regulated genes with the highest p value among the molecular functions and cellular components is interrelated to Disc development (GO: 0035218), ECM structural components (GO: 0005201) and extracellular regions (GO: 0005576) (Figure 3A-3C). In addition, we have constructed a miRNA-mRNA network map through Cytoscape software (Figure 3D). To further probe the potential targets of miR-31-5p, we compiled all the predicted genes into a Venn analysis map (Figure 3E). According to the result, we demonstrated that the SDF-1/CXCR7 axis is the target of miR-31-5p (Figure 3F). Besides, miR-31-5p is proven to be highly conserved among species (Figure 3G). To further verify the association between the SDF-1/CXCR7 axis and miR-31-5p, luciferase reporter gene analysis was employed to test the association between them. The results demonstrated that the relative luciferase reporter activity of wild type (WT) co-transfected with miR-31-5p mimic in primary human NP cells was meaningfully lower than that of mutant (mut) cells transfected with miR-31-5p mimic (Figure 3H, *P* < 0.001). We further verified this result at the protein level. The results of western blot demonstrated that the level of SDF-1 protein in miR-31-5p mimic group declined (Figure 3I). The above results indicate that the SDF-1/CXCR7 axis is the target of miR-31-5p.

**Figure.3.**
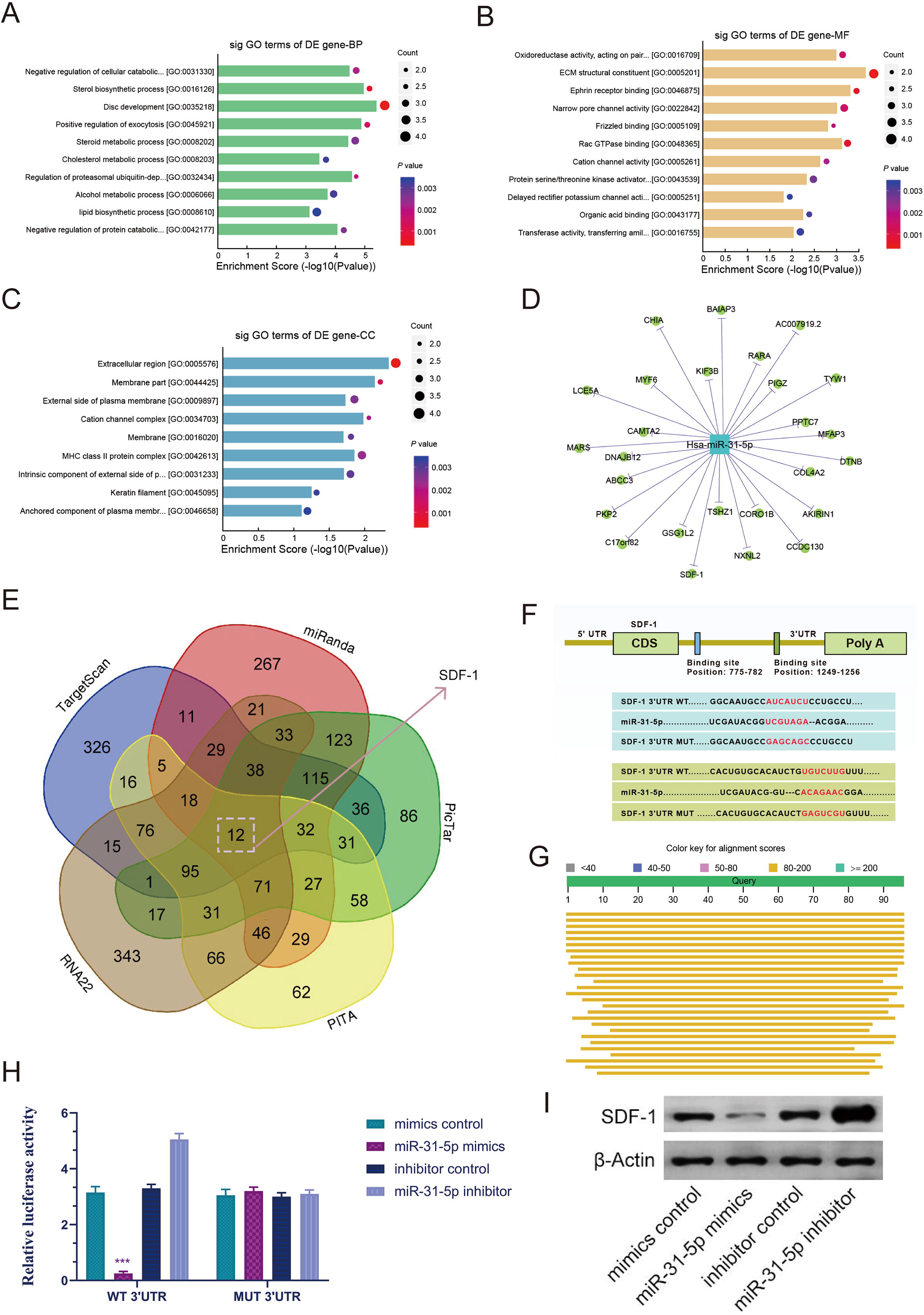
The relevance of miR-31-5p to SDF-1/CXCR7 axis. (A, B, C) For biological processes, molecular functions and cellular components, the down-regulated GO term with the highest p value. (D) The target of miR-31-5p was verified by Cytoscape. (C) The Venn diagram predicts that SDF-1/CXCR7 axis is the target of miR-31-5p. (E) Venn diagram displaying miR-31-5p computationally predicted to target SDF-1 by different algorithms. (F) The mRNA 3’UTR of SDF-1 and the putative miR-31-5p binding site sequence has high sequence conservation and complementarity with miR-31-5p. (G) miR-31-5p is highly conservative. (H) The wild-type or mutant SDF-1 3’UTR reporter plasmid and miR-31-5p mimic or inhibitor are co-transfected into human NP cells. (I) The expression level of SDF-1 was valued through western blot. n = 3. IDD, Intervertebral disc degeneration; miR: microRNA; NP: Nucleus pulposus; GO: gene ontology; mut: mutant; WT: wild type; SDF-1: stromal cell-derived Faceor-1; CXCR: C-X-C chemokine receptor. ^***^*P <* 0.001.

### miR-31-5p level regulated IDD via SDF-1/CXCR7 axis

SDF-1/CXCR7 is closely interrelated to the occurrence of many diseases, and previous studies have demonstrated that SDF-1/CXCR7 is interrelated to the occurrence of IDD [36–38]. As shown in Figure 4A, the SDF-1/CXCR7 signaling pathway is significantly rich in genes and genomic pathways of the Kyoto Protocol. Through further research, we confirmed that miR-31-5p regulates IDD through the SDF-1/CXCR7 axis pathway. We transfected the cultured primary human NP cells with miR-31-5p mimic, miR-31-5p inhibitor or its negative control. Western blot results demonstrated that in NP cells, the protein levels of SDF-1, CXCR7, ADAMTS-5 and MMP13 of miR-31-5p mimics declined, while in NP cells transfected with miR-31-5p inhibitor, SDF-1, CXCR7, ADAMTS-5 and MMP13 protein levels increased. In addition, the effects of SDF-1 small interfering RNA (siRNA) on SDF-1, CXCR7, ADAMTS-5 and MMP13 are similar to those induced by miR-31-5p mimics. This manifests that miR-31-5p regulates IDD through the SDF-1/CXCR7 axis pathway (Figure 4B). Further experiments were executed to verify the association between miR-31-5p and SDF-1/CXCR7 axis (Figure 4C and 4D). The above experimental results show that miR-31-5p acts through the SDF-1/CXCR7 axis pathway.

**Figure.4.**
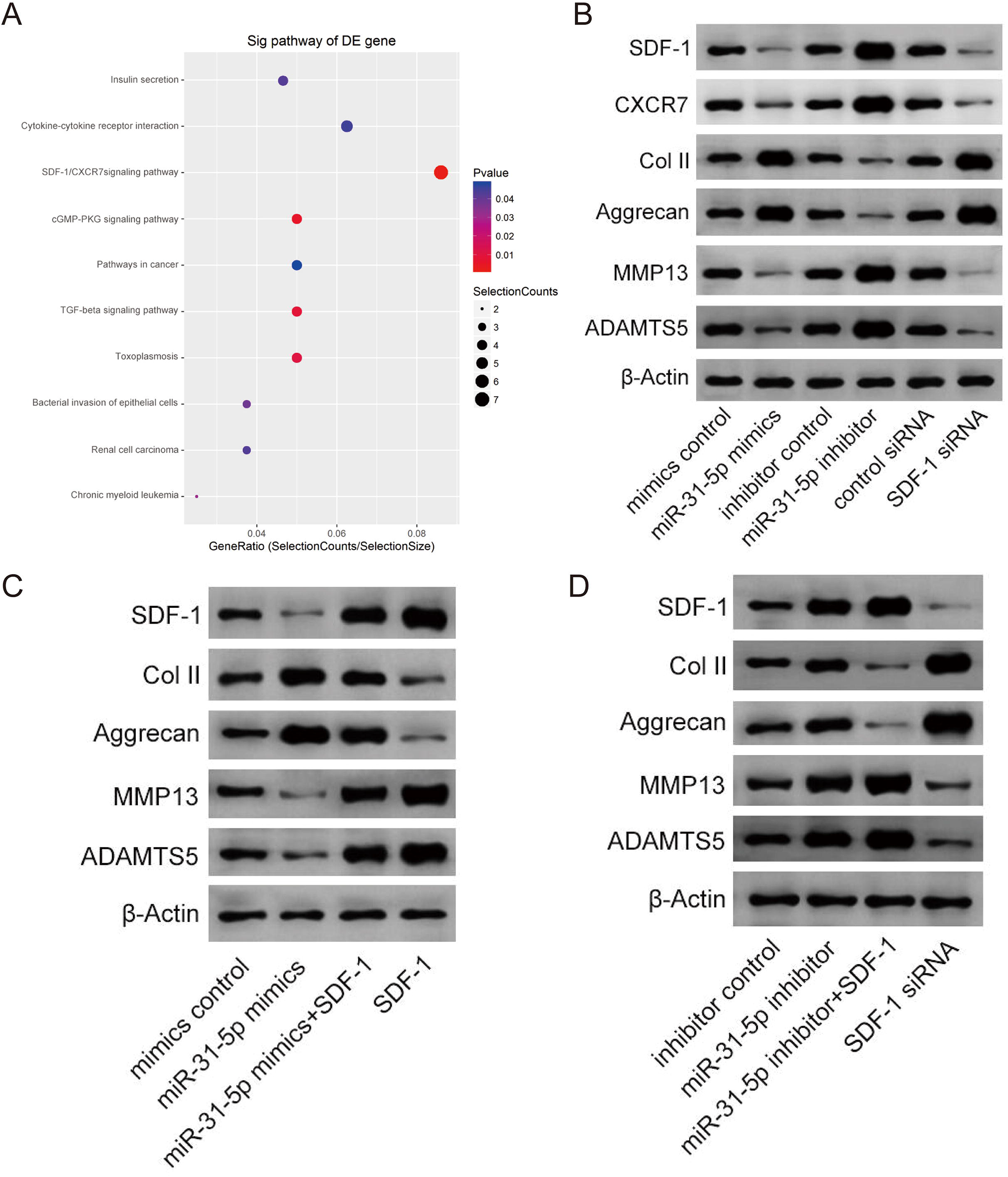
miR-31-5p expression regulates IDD via SDF-1/CXCR7 axis. (A) The IDD-rich SDF-1/CXCR7 pathway was analyzed by KEGG. (B) The expression level of SDF-1, CXCR7, Col II, Aggrecan, MMP13 and ADAMT5 was valued through western blot. (C) The expression level of SDF-1, Col II, Aggrecan, MMP13 and ADAMT5 was valued through western blot. (D) The expression level of SDF-1, Col II, Aggrecan, MMP13 and ADAMT5 was valued through western blot. IDD, Intervertebral disc degeneration; miR: microRNA; Col II: Type II collagen; MMP: matrix metalloprotein; ADAMTS: a disintegrin-like and metalloproteinase with thrombospondin motifs; KEGG, Kyoto Encyclopedia of Genes and Genomes; SDF-1: stromal cell-derived Faceor-1; CXCR: C-X-C chemokine receptor.

### Upregulation of miR-31-5p level prevented IDD development

We further studied the part of miR-31-5p in IDD and the molecular mechanisms involved. We induced the IDD model by WT mice, and then injected miR-31-5p mimic or inhibitor NP and control NP locally on 1, 7, and 14 days after surgery (Figure 5A). We monitored the in vivo targeting ability of NP in real time. The results revealed that miR-31-5p mediated by NPs revealed a good delivery function in mice (Figure 5B). In order to further probe the part of miR-31-5p in IDD, we conducted further tests through radiography and histological evaluation. The results revealed that compared with the control group, the local delivery of miR-31-5p mimic NPs significantly protected the IVD structure, which indicated that miR-31-5p overexpression had a protective function on the surgically induced IDD model (Figure 5C-5F). NPs treated with miR-31-5p mimic significantly reduced the level of MMP13, while the level of col II increased. The miR-31-5p inhibitor group had the opposite effect (Figure 5G). We tested the apoptosis of NP cells after different treatments, and the results of TUNEL staining revealed that NP cell apoptosis was significant in mice treated with miR-31-5p mimic NPs. Reduce (Figure 5H). The above results show that overexpression of miR-31-5p has a significant effect on the treatment of IDD, which indicates that miR-31-5p is a potential therapeutic target for IDD.

**Figure.5.**
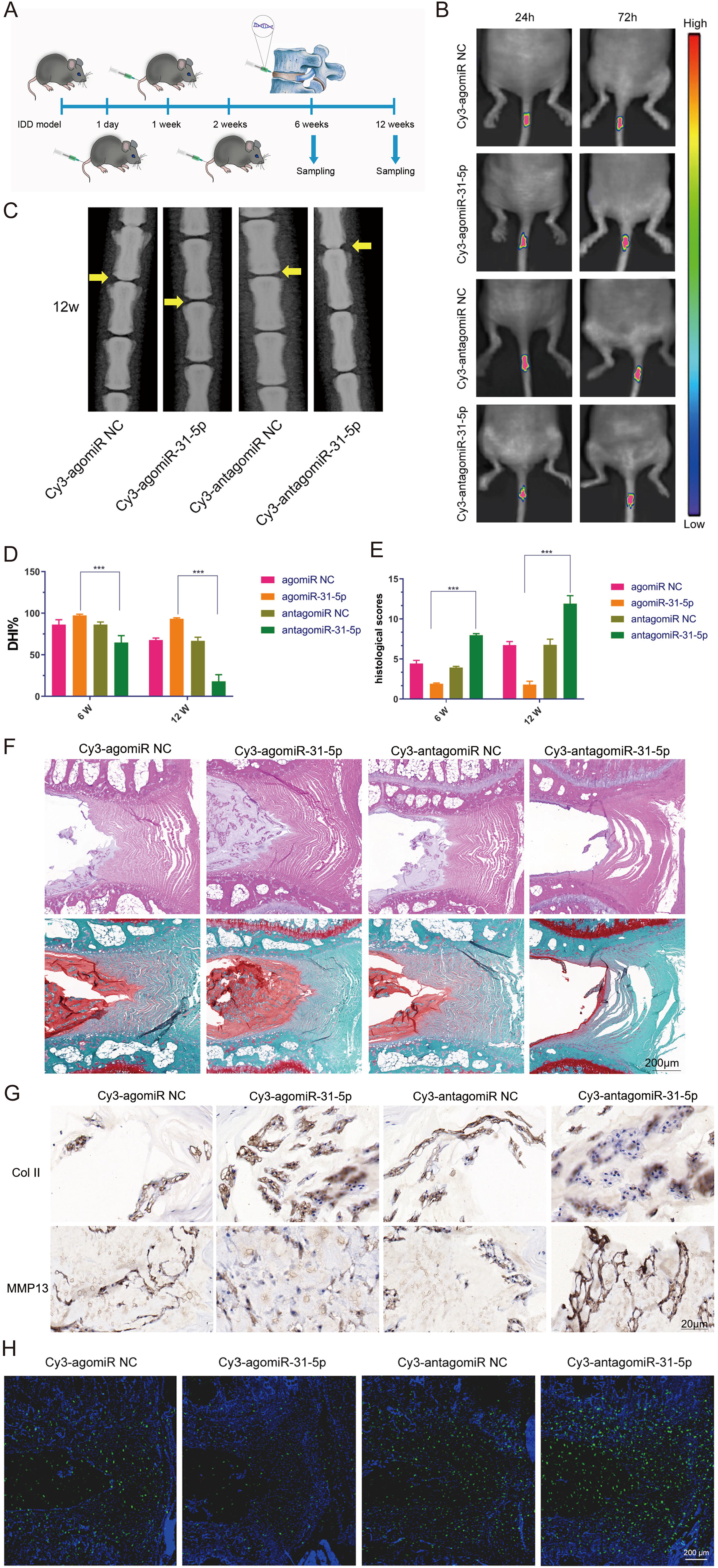
Upregulation of miR-31-5p expression prevented IDD development. (A) Experiments in which miR-31-5p mimics, miR-31-5p inhibitors or their negative controls were injected at 1, 1 and 2 weeks after surgery. (B) Time-dependent fluorescence images of mice treated with Cy3-miR-31-5p for 24h and 72h. From blue to red represents the change in fluorescence signal intensity from low to high. (C) X-ray evaluation of intervertebral disc degeneration. (D-F) The DHI of 6W and 12W under different treatments and the changes in histology were disclosed. (G) Immunostaining of Col II and MMP13 in IDD model treated with miR-31-5p NP. Scale bar = 20μm. (H) TUNEL staining was executed on the intervertebral disc to determine the apoptotic activity, scale bar = 200μm. IDD, Intervertebral disc degeneration; NC: Negative control; miR: microRNA; Col II: Type II collagen; MMP: matrix metalloprotein. \*\*\**P <* 0.001.

## Discussion

Many studies have manifested that NP cells are essential for maintaining the structural integrity of intervertebral discs, and the phenotype of NP cells is closely interrelated to the pathogenesis of IDD [39, 40]. Regarding the current treatment methods and methods, in-depth research is needed to promote the improvement of prevention and treatment methods. Recent studies have reported that certain specific miRNAs have been identified as important regulatory factors and biomarkers of IDD, which are important substances for the prevention and treatment of IDD [41, 42]. The underlying mechanism is to regulate certain chemokines and transcription and translation in cells through specific miRNAs, and then regulate the cell phenotype [17, 43, 44].

Microarray detection of IDD NP was executed to analyze the differential miRNA level profile between IDD and normal tissues. miR-31-5p is closely interrelated to the occurrence and development of many diseases [45–47]. In this study, the part of miR-31-5p in the occurrence and development of IDD was further verified. In order to further probe the imbalance of miR-31-5p in IDD, the level of miR-31-5p was disclosed by qRT-PCR, and it was demonstrated that miR-31-5p was down-regulated in IDD. Further FISH maps and methylation conditions also verified this result. In order to clarify the function of miR-31-5p level on NP cells, the function of miR-31-5p level and silencing miR-31-5p on the phenotype of NP cells was studied by regulating the level of miR-31-5p. The results proved that overexpression of miR-31-5p is interrelated to increased proliferation of NP cells, inhibition of apoptosis, increased ECM formation, and inhibition of matrix degrading enzymes. These phenotypic changes are the key biological process of IDD. The increase in ECM synthesis is manifested by the increase in the ingredients of Col II and Aggrecan, while the decrease of the ingredients in ADAMTS-5 and MMP13 [48, 49]. Therefore, miR-31-5p is involved in the pathogenesis of IDD.

In order to further study how miR-31-5p regulates IDD, GO analysis of dysregulated mRNA was carried out, miRNA-mRNA network diagram was constructed, Venn analysis and dual luciferase analysis demonstrated that SDF-1/CXCR7 axis is the target of miR-31-5p. Previous studies have manifested that SDF-1/CXCR7 is involved in the regulation of many diseases, such as acute myocardial infarction (AMI), degenerative disc disease (DDD), steroid-induced osteonecrosis of the femoral head (SONFH) [36, 50, 51]. Prior to this, many studies have demonstrated that there is a targeting association between SDF-1/CXCR7 and miRNA ^[52, 53]^. The results of this study support that the increase in miR-31-5p level is consistent with the decrease in SDF-1 protein level through Western blot detection.

It is further verified that miR-31-5p regulates the changes of NP phenotype through the SDF-1/CXCR7 axis pathway. It is consistent with that in NP cells, the overexpression of miR-31-5p is consistent with the reduction of protein levels of SDF-1, CXCR7, ADAMTS-5 and MMP13. Silencing miR-31-5p is consistent with increased protein levels of SDF-1, CXCR7, ADAMTS-5 and MMP13. The effects of SDF-1 small interfering RNA (siRNA) on SDF-1, CXCR7, ADAMTS-5 and MMP13 are similar to those induced by miR-31-5p mimics. This result also indicates that there is a negative regulatory association between miR-31-5p and SDF-1/CXCR7 axis. SDF-1 can offset the function of overexpression of miR-31-5p on the protein levels of SDF-1, col II, Aggrecan, ADAMTS-5 and MMP13. It is worth noting that the function of SDF-1 on miR-31-5p inhibition is limited to col II and Aggrecan. The above information manifests that miR-31-5p regulates IDD through the SDF-1/CXCR7 axis pathway. Although other study had reported some critical biological functions of miR-31-5p, this was the first time to reveal its function on SDF1/CXCR7 axis.

The important clinical significance of miR-31-5p and its downstream pathways was further confirmed by in vivo examination using an inducible IDD animal model. As expected, in terms of protecting the phenotype of NP cells, overexpression of miR-31-5p effectively alleviated the symptoms of IDD. Therefore, as an effective regulator of IDD, miR-31-5p has therapeutic potential. It is not the first report that miRNAs play a positive role in the treatment of IDD. Many previous studies have manifested that miRNAs regulate IDD through β-catenin, MMP, apoptosis and cell proliferation [54–60]. It is worth emphasizing that the level of miRNAs may be regulated by the methylation of the promoter region, which may affect certain phenotypes [61, 62]. In this research, according to the prediction of CpG islands in the promoter, three strong CpG islands were identified in miR-31-5p interrelated promoters. This makes us hypothesize that miR-31-5p in its host gene may be regulated by promoter methylation. Further experiments proved that hypermethylation may contribute to the loss of miR-31-5p in NP cells. In addition, a comprehensive understanding of the relevant molecular mechanisms will have an function on the efficacy of treatment.

In addition to the important findings of this study, restrictions on the scope of the study still exist. First, although the function of down-regulation of miR-31-5p on the phenotype of NP cells has been demonstrated, the mechanism of down-regulation of miR-31-5p is not fully understood. Secondly, other potential effects of manually increasing miR-31-5p are not yet known. These issues may be further studied in follow-up research work.

In conclusion, miR-31-5p reduces IDD by targeting SDF-1/CXCR7 to regulate cell proliferation, apoptosis, ECM and matrix degradation. These findings lay the foundation for follow-up research and the understanding and understanding of IDD, and at the same time provide a promising therapeutic target for IDD treatment.

## Conclusions

In this study, our results indicate that miR-31-5p has a potential part in the proliferation, apoptosis, ECM formation of NPs, and matrix degrading enzymes in NP cells. These effects may be achieved through the SDF-1/CXCR7 axis, and further in vitro experiments have further verified the part of miR-31-5p. Importantly, our results provide evidence for miR-31-5p as a potential target, diagnostic indicator and prognostic indicator for IDD patients.

## References

1. Murray CJ and Lopez AD. Measuring the global burden of disease. The New England journal of medicine. 2013; 369(5):448–457.

2. Richardson SM, Kalamegam G, Pushparaj PN, Matta C, Memic A, Khademhosseini A, Mobasheri R, Poletti FL, Hoyland JA and Mobasheri A. Mesenchymal stem cells in regenerative medicine: Focus on articular cartilage and intervertebral disc regeneration. Methods. 2016; 99:69–80.

3. Adams MA and Roughley PJ. What is intervertebral disc degeneration, and what causes it? Spine. 2006; 31(18):2151–2161.

4. Palepu V, Kodigudla M and Goel VK. Biomechanics of disc degeneration. Advances in orthopedics. 2012; 2012:726210.

5. Ghannam M, Jumah F, Mansour S, Samara A, Alkhdour S, Alzuabi MA, Aker L, Adeeb N, Massengale J and Oskouian RJ. Surgical anatomy, radiological features, and molecular biology of the lumbar intervertebral discs. J Clinical Anatomy. 2017; 30(2):251–266.

6. Nies DE, Hemesath TJ, Kim JH, Gulcher JR and Stefansson K. The complete cDNA sequence of human hexabrachion (Tenascin). A multidomain protein containing unique epidermal growth factor repeats. The Journal of biological chemistry. 1991; 266(5):2818–2823.

7. Cui S and Zhang L. circ_001653 Silencing Promotes the Proliferation and ECM Synthesis of NPCs in IDD by Downregulating miR-486-3p-Mediated CEMIP. Molecular therapy Nucleic acids. 2020; 20:385–399.

8. Tan H, Zhao L, Song R, Liu Y and Wang L. microRNA‐665 promotes the proliferation and matrix degradation of nucleus pulposus through targeting GDF5 in intervertebral disc degeneration. Journal of cellular biochemistry. 2018; 119(9):7218–7225.

9. Bartel DP. MicroRNAs: genomics, biogenesis, mechanism, and function. Cell. 2004; 116(2):281–297.

10. Hammond SM. An overview of microRNAs. Advanced drug delivery reviews. 2015; 87:3–14.

11. Lei SF, Papasian CJ and Deng HW. Polymorphisms in predicted miRNA binding sites and osteoporosis. Journal of Bone Miner Res. 2011; 26(1):72–78.

12. Kolhe R, Hunter M, Liu S, Jadeja RN, Pundkar C, Mondal AK, Mendhe B, Drewry M, Rojiani MV and Liu Y. Gender-specific differential expression of exosomal miRNA in synovial fluid of patients with osteoarthritis. Scientific reports. 2017; 7(1):1–14.

13. Nakasa T, Miyaki S, Okubo A, Hashimoto M, Nishida K, Ochi M and Asahara H. Expression of microRNA‐146 in rheumatoid arthritis synovial tissue. Arthritis Rheumatism. 2008; 58(5):1284–1292.

14. Ji M-l, Zhang X-j, Shi P-l, Lu J, Wang S-z, Chang Q, Chen H and Wang C. Downregulation of microRNA-193a-3p is involved in invertebral disc degeneration by targeting MMP14. Journal of molecular medicine. 2016; 94(4):457–468.

15. Song Y-Q, Karasugi T, Cheung KM, Chiba K, Ho DW, Miyake A, Kao PY, Sze KL, Yee A and Takahashi A. Lumbar disc degeneration is linked to a carbohydrate sulfotransferase 3 variant. The Journal of clinical investigation. 2013; 123(11):4909–4917.

16. Ji Ml, Lu J, Shi Pl, Zhang Xj, Wang Sz, Chang Q, Chen H and Wang C. Dysregulated miR‐98 contributes to extracellular matrix degradation by targeting IL‐6/STAT3 signaling pathway in human intervertebral disc degeneration. Journal of Bone Mineral Research. 2016; 31(4):900–909.

17. Wang C, Wang WJ, Yan YG, Xiang YX, Zhang J, Tang ZH and Jiang ZS. MicroRNAs: New players in intervertebral disc degeneration. Clinica chimica acta; international journal of clinical chemistry. 2015; 450:333–341.

18. Yu T, Ma P, Wu D, Shu Y and Gao W. Functions and mechanisms of microRNA-31 in human cancers. Biomedicine & pharmacotherapy = Biomedecine & pharmacotherapie. 2018; 108:1162–1169.

19. Wang S, Hu J, Zhang D, Li J, Fei Q and Sun Y. Prognostic role of microRNA-31 in various cancers: a meta-analysis. Tumor Biology. 2014; 35(11):11639–11645.

20. Wang X-Q, Tu W-Z, Guo J-B, Song G, Zhang J, Chen C-C and Chen P-J. A bioinformatic analysis of microRNAs’ role in human intervertebral disc degeneration. Pain medicine. 2019; 20(12):2459–2471.

21. Wang X, Zhang Y, Jiang B, Zhang Q, Zhou R, Zhang L and Wang C. Study on the role of Hsa-miR-31-5p in hypertrophic scar formation and the mechanism. Experimental cell research. 2017; 361(2):201–209.

22. Ji ML, Zhang XJ, Shi PL, Lu J, Wang SZ, Chang Q, Chen H and Wang C. Downregulation of microRNA-193a-3p is involved in invertebral disc degeneration by targeting MMP14. Journal of Molecular Medicine. 2015; 94(4):457–468.

23. Xia X, Guo J, Lu F and Jiang J. SIRT1 Plays a Protective Role in Intervertebral Disc Degeneration in a Puncture-induced Rodent Model. Spine. 2015; 40(9):E515–524.

24. O’Connell GD, Vresilovic EJ and Elliott DM. Comparison of animals used in disc research to human lumbar disc geometry. Spine. 2007; 32(3):328–333.

25. Schaer TP, Vresilovic EJ, Mechanics A, Reynolds RS, Square K, Elliott DM, Showalter BL, Beckstein JC, Martin JT and Beattie EE. Comparison of animal discs used in disc research to human lumbar disc: torsion mechanics and collagen content. Spine. 2012; 37(15):E900.

26. Jesse, Beckstein, Sounok, Sen, Thomas, Schaer, Edward J. Comparison of animal discs used in disc research to human lumbar disc: axial compression mechanics and glycosaminoglycan content. Spine. 2008.

27. Han B, Zhu K, Li FC, Xiao YX, Feng J, Shi ZL, Lin M, Wang J and Chen QX. A simple disc degeneration model induced by percutaneous needle puncture in the rat tail. Spine. 2008; 33(18):1925–1934.

28. Rousseau MA, Ulrich JA, Bass EC, Rodriguez AG, Liu JJ and Lotz JC. Stab incision for inducing intervertebral disc degeneration in the rat. Spine. 2007; 32(1):17–24.

29. Masuda K, Aota Y, Muehleman C, Imai Y, Okuma M, Thonar EJ, Andersson GB and An HS. A novel rabbit model of mild, reproducible disc degeneration by an anulus needle puncture: correlation between the degree of disc injury and radiological and histological appearances of disc degeneration. Spine. 2005; 30(1):5–14.

30. Lipson SJ and Muir H. Experimental intervertebral disc degeneration: morphologic and proteoglycan changes over time. Arthritis and rheumatism. 2010; 24(1):12–21.

31. Tam V, Chan WCW, Leung VYL, Cheah KSE, Cheung KMC, Sakai D, McCann MR, Bedore J, Seguin CA and Chan D. Histological and reference system for the analysis of mouse intervertebral disc. Journal of orthopaedic research : official publication of the Orthopaedic Research Society. 2018; 36(1):233–243.

32. Boos N, Weissbach S, Rohrbach H, Weiler C and Nerlich AG. Classification of age-related changes in lumbar intervertebral discs: 2002 Volvo Award in basic science. Spine. 2003; 27(23):2631–2644.

33. Horner HA, Roberts S, Bielby RC, Menage J, Evans H and Urban JP. Cells from different regions of the intervertebral disc: effect of culture system on matrix expression and cell phenotype. Spine. 2002; 27(10):1018–1028.

34. Han, Qadir, X. V and Zhang. MiR-185 inhibits hepatocellular carcinoma growth by targeting the DNMT1/PTEN/Akt pathway. American Journal of Pathology Official Publication of the American Association of Pathologists. 2014.

35. Yang Y, Zhong Z, Zhao Y, Ren K and Li N. LincRNA-SLC20A1 (SLC20A1) promotes extracellular matrix degradation in nucleus pulposus cells in human intervertebral disc degeneration by targeting the miR-31-5p/MMP3 axis. Int J Clin Exp Pathol. 2019; 12(9):3632–3643.

36. Zhang H, Wang P, Zhang X, Zhao W, Ren H and Hu Z. SDF1/CXCR7 Signaling Axis Participates in Angiogenesis in Degenerated Discs via the PI3K/AKT Pathway. DNA and cell biology. 2019; 38(5):457–467.

37. Luo Y, Azad AK, Karanika S, Basourakos SP, Zuo X, Wang J, Yang L, Yang G, Korentzelos D and Yin J. Enzalutamide and CXCR7 inhibitor combination treatment suppresses cell growth and angiogenic signaling in castration‐resistant prostate cancer models. International journal of cancer. 2018; 142(10):2163–2174.

38. Hain S. (2019). The Roles of CXCR4 and CXCR7 in Melanocyte and Melanoma Motility. California State University, Northridge).

39. Chen J, Xie JJ, Jin MY, Gu YT, Wu CC, Guo WJ, Yan YZ, Zhang ZJ, Wang JL, Zhang XL, Lin Y, Sun JL, Zhu GH, et al. Sirt6 overexpression suppresses senescence and apoptosis of nucleus pulposus cells by inducing autophagy in a model of intervertebral disc degeneration. Cell Death Dis. 2018; 9(2):56.

40. Jiang L, Zhang X, Zheng X, Ru A, Ni X, Wu Y, Tian N, Huang Y, Xue E, Wang X and Xu H. Apoptosis, senescence, and autophagy in rat nucleus pulposus cells: Implications for diabetic intervertebral disc degeneration. Journal of orthopaedic research : official publication of the Orthopaedic Research Society. 2013; 31(5):692–702.

41. Xu YQ, Zhang ZH, Zheng YF and Feng SQ. Dysregulated miR-133a Mediates Loss of Type II Collagen by Directly Targeting Matrix Metalloproteinase 9 (MMP9) in Human Intervertebral Disc Degeneration. Spine. 2016; 41(12):E717–724.

42. Cai P, Yang T, Jiang X, Zheng M, Xu G and Xia J. Role of miR-15a in intervertebral disc degeneration through targeting MAP3K9. Biomedicine Pharmacotherapy. 2017; 87:568–574.

43. Ji ML, Jiang H, Zhang XJ, Shi PL, Li C, Wu H, Wu XT, Wang YT, Wang C and Lu J. Preclinical development of a microRNA-based therapy for intervertebral disc degeneration. Nature communications. 2018; 9(1):5051.

44. Sherafatian M, Abdollahpour HR, Ghaffarpasand F, Yaghmaei S, Azadegan M and Heidari M. MicroRNA expression profiles, target genes, and pathways in intervertebral disk degeneration: a meta-analysis of 3 microarray studies. World neurosurgery. 2019; 126:389–397.

45. Chen G, Han Y, Feng Y, Wang A, Li X, Deng S, Zhang L, Xiao J, Li Y and Li N. Extract of Ilex rotunda Thunb alleviates experimental colitis-associated cancer via suppressing inflammation-induced miR-31-5p/YAP overexpression. Phytomedicine : international journal of phytotherapy and phytopharmacology. 2019; 62:152941.

46. Gupta P, Yadav RP, Rawat P and Baranwal S. Commentary on: MicroRNA-31 Reduces Inflammatory Signaling and Promotes Regeneration in Colon Epithelium, and Delivery of Mimics in Microspheres Reduces Colitis in Mice. DOI: 10.1053/j.gastro.2019.02.023. PMID: 30779922. Frontiers in Immunology. 2019; 10:2649.

47. He J, He J, Min L, He Y, Guan H, Wang J and Peng X. Extracellular vesicles transmitted miR‐31‐5p promotes sorafenib resistance by targeting MLH1 in renal cell carcinoma. International Journal of Cancer. 2020; 146(4):1052–1063.

48. Takaishi H, Kimura T, Dalal S, Okada Y and D’Armiento J. Joint diseases and matrix metalloproteinases: a role for MMP-13. Current pharmaceutical biotechnology. 2008; 9(1):47–54.

49. Gendron C, Kashiwagi M, Lim NH, Enghild JJ, Thogersen IB, Hughes C, Caterson B and Nagase H. Proteolytic activities of human ADAMTS-5: comparative studies with ADAMTS-4. The Journal of biological chemistry. 2007; 282(25):18294–18306.

50. Zhang S, Yue J, Ge Z, Xie Y, Zhang M and Jiang L. Activation of CXCR7 alleviates cardiac insufficiency after myocardial infarction by promoting angiogenesis and reducing apoptosis. Biomedicine & pharmacotherapy = Biomedecine & pharmacotherapie. 2020; 127:110168.

51. Kong L, Zuo R, Wang M, Wang W, Xu J, Chai Y, Guan J and Kang Q. Silencing MicroRNA-137-3p, which Targets RUNX2 and CXCL12 Prevents Steroid-induced Osteonecrosis of the Femoral Head by Facilitating Osteogenesis and Angiogenesis. International journal of biological sciences. 2020; 16(4):655.

52. Shan C and Ma Y. MicroRNA-126/stromal cell-derived factor 1/CXC chemokine receptor type 7 signaling pathway promotes post-stroke angiogenesis of endothelial progenitor cell transplantation. Molecular medicine reports. 2018; 17(4):5300–5305.

53. Yang L-X, Wei C-L, Guo M-L, Zhang Y, Bai F and Ma S-G. Improvement of therapeutic effects of mesenchymal stem cells in myocardial infarction through genetic suppression of microRNA-142. Oncotarget. 2017; 8(49):85549.

54. Feng G, Zha Z, Huang Y, Li J, Wang Y, Ke W, Chen H, Liu L, Song Y and Ge Z. Sustained and Bioresponsive Two‐Stage Delivery of Therapeutic miRNA via Polyplex Micelle‐Loaded Injectable Hydrogels for Inhibition of Intervertebral Disc Fibrosis. Advanced healthcare materials. 2018; 7(21):1800623.

55. Ge C and Li C. Expression and regulatory role of miRNA-222 in intervertebral disc degeneration (IDD). Biotechnology Biotechnological Equipment. 2019; 33(1):1553–1559.

56. Wei Y, Nazari-Jahantigh M, Neth P, Weber C and Schober A. MicroRNA-126,-145, and-155: a therapeutic triad in atherosclerosis? Arteriosclerosis, thrombosis, vascular biology. 2013; 33(3):449–454.

57. Wang HQ, Yu XD, Liu ZH, Cheng X, Samartzis D, Jia LT, Wu SX, Huang J, Chen J and Luo ZJ. Deregulated miR‐155 promotes Fas‐mediated apoptosis in human intervertebral disc degeneration by targeting FADD and caspase‐3. The Journal of pathology. 2011; 225(2):232–242.

58. Liu G, Cao P, Chen H, Yuan W, Wang J and Tang X. MiR-27a regulates apoptosis in nucleus pulposus cells by targeting PI3K. PloS one. 2013; 8(9):e75251.

59. Li Z, Shen J, Wu WK, Yu X, Liang J, Qiu G and Liu J. Leptin induces cyclin D1 expression and proliferation of human nucleus pulposus cells via JAK/STAT, PI3K/Akt and MEK/ERK pathways. PloS one. 2012; 7(12):e53176.

60. Yu X, Li Z, Shen J, Wu WK, Liang J, Weng X and Qiu G. MicroRNA-10b promotes nucleus pulposus cell proliferation through RhoC-Akt pathway by targeting HOXD10 in intervetebral disc degeneration. PloS one. 2013; 8(12):e83080.

61. Augoff K, McCue B, Plow EF and Sossey-Alaoui K. miR-31 and its host gene lncRNA LOC554202 are regulated by promoter hypermethylation in triple-negative breast cancer. Mol Cancer. 2012; 11(1):5.

62. Li J, Zhang S, Zou Y, Wu L, Pei M and Jiang Y. miR‐145 promotes miR‐133b expression through c‐myc and DNMT3A‐mediated methylation in ovarian cancer cells. Journal of cellular physiology. 2020; 235(5):4291–4301.

